# GOLVEN peptides regulate lateral root spacing as part of a negative feedback loop on the establishment of auxin maxima

**DOI:** 10.1101/2022.09.26.509595

**Authors:** Joris Jourquin, Ana Ibis Fernandez, Ke Xu, Jan Šimura, Karin Ljung, Tom Beeckman

**Author notes:** Correspondence /.

## Abstract

The initiation of lateral roots in *Arabidopsis* requires the accumulation of auxin in lateral root founder cells, yielding a local auxin maximum. The positioning of these auxin maxima along the primary root determines the density and spacing of lateral roots. The GOLVEN6 (GLV6) and GLV10 signaling peptides and their receptors have been established as regulators of lateral root spacing via an inhibitory effect on lateral root initiation. However, it remained unclear how these GLV peptides interfere with auxin signaling or homeostasis. Here we show that GLV6/10 signaling regulates the expression of a subset of auxin response genes during lateral root initiation, downstream of the canonical auxin signaling pathway, while simultaneously inhibiting the establishment of auxin maxima in xylem-pole pericycle cells. We present genetic evidence that this inhibitory effect relies on the activity of the PIN3 and PIN7 auxin export proteins. Furthermore, GLV6/10 peptide signaling was found to enhance PIN7 abundance in the plasma membrane of xylem-pole pericycle cells, which likely stimulates auxin efflux from these cells. Based on these findings, we propose a model in which the GLV6/10 signaling pathway serves as a negative feedback loop that contributes to the robust patterning of auxin maxima along the primary root.

## Introduction

Roots enable plants to take up water and nutrients from the soil, interact with soil microbiota, and fix themselves to the substrate. Hence, a flexible root system, with the ability to branch out, is essential for the optimization of the root surface area, the exploration of different micro-environments in the soil, and the formation of a more extensive support structure. Control over the architecture of the root system is therefore of particular importance for plants, as illustrated by their ability to adjust the density and spacing of lateral roots (LRs) along the primary root based on the availability of water and nutrients, as well as the presence of certain microorganisms in the rhizosphere (López-Bucio *et al*., 2002, 2007; Shin *et al*., 2007; Zamioudis *et al*., 2013; Bao *et al*., 2014; Orman-Ligeza *et al*., 2018; Orosa-Puente *et al*., 2018; Kong *et al*., 2020). Accordingly, a better understanding of the mechanisms that determine when and where LR development is initiated might enable us to make beneficial modifications to the root system architecture of certain crops through breeding efforts or biotechnological approaches. The chief regulator of LR development is the phytohormone auxin (Lavenus *et al*., 2013; Du and Scheres, 2018). In *Arabidopsis thaliana*, LRs arise from pairs of abutting xylem-pole pericycle (XPP) cells, called lateral root founder cells (LRFCs) (Dubrovsky *et al*., 2000, 2008; Beeckman *et al*., 2001; Parizot *et al*., 2008). In the young maturation zone, auxin accumulates in these cells, which can be visualized using the auxin-responsive *DR5* promoter (Ulmasov *et al*., 1997). DR5*-* marked patches of XPP cells are therefore often referred to as pre-branch sites (van Norman *et al*., 2013). Upon lateral root initiation, LRFCs undergo an asymmetric anticlinal cell division that is essential for further lateral root development (Malamy & Benfey, 1997; De Smet et al., 2007; De Rybel et al., 2010; Goh et al., 2012a). Extensive research has shown that the local accumulation of auxin in LRFCs and an intact auxin signaling pathway are instrumental for proper LR initiation and development. Inside the LRFCs, auxin induces the breakdown of AUX/IAA proteins via TRANSPORT INHIBITOR RESPONSE 1 (TIR1) and TIR1-like AUXIN SIGNALING F-BOX PROTEINs (AFBs) (Kepinski and Leyser, 2005; Parry *et al*., 2009), which releases AUXIN RESPONSE FACTOR (ARF) transcription factors to modulate the expression of their target genes (Ulmasov *et al*., 1997, 1999; Gray *et al*., 2001; Tiwari *et al*., 2001). Several AUX/IAAs and ARFs are known to be involved in lateral root development, but IAA14/SLR, ARF7 and ARF19 seem to play the most prominent role during lateral root initiation (Fukaki *et al*., 2002; Okushima *et al*., 2005; Goh *et al*., 2012*b*). Via this pathway, auxin modulates the expression of a host of auxin response genes that orchestrate the initiation and development of lateral roots (Okushima *et al*., 2005; Vanneste *et al*., 2005). The local auxin accumulation in LRFCs required for the induction of this pathway is achieved through careful regulation of its biosynthesis, storage and degradation (Nakazawa *et al*., 2001; Rampey *et al*., 2004; Ljung *et al*., 2005; Porco *et al*., 2016; Zhang *et al*., 2016; Casanova-Sáez *et al*., 2022), combined with directional auxin transport, which is mediated by 3 main classes of transmembrane transporters; the AUXIN RESISTANT 1 (AUX1) and LIKE-AUX1 (LAX) importers (Swarup and Péret, 2012), the ATP-BINDING CASSETTE TYPE-B (ABCB) exporters (Geisler *et al*., 2017), and the PIN-FORMED (PIN) exporters (Adamowski and Friml, 2015). Defects in the auxin transport machinery can affect the spacing and density of lateral roots, and chemical inhibition of polar auxin transport can even block lateral root initiation altogether, highlighting the importance of local auxin accumulation in LRFCs (Casimiro, 2001; Marchant *et al*., 2002; Benková *et al*., 2003; Laskowski *et al*., 2008; Mravec *et al*., 2008; Swarup *et al*., 2008; Marhavý *et al*., 2013).

In addition to auxin, several small secreted signaling peptides and their receptors have also been shown to be involved in the regulation of LR initiation and spacing (De Smet et al., 2008; Araya et al., 2014; Cho et al., 2014; Czyzewicz et al., 2015; Murphy et al., 2016; Roberts et al., 2016; Dimitrov and Tax, 2018; Dong et al., 2019; Toyokura et al., 2019; Fernandez et al., 2020; Jourquin et al., 2022). These include GOLVEN (GLV) 6 and GLV10, two members of the GOLVEN/ROOT GROWTH FACTOR/CLE-LIKE peptide family, and the receptor-like kinases (RLKs) responsible for their perception, called ROOT GROWTH FACTOR INSENSITIVE (RGI) 1-5 (Fernandez *et al*., 2013, 2015, 2020; Ou *et al*., 2016; Song *et al*., 2016). *GLV6* and *GLV10* are auxin-inducible genes that start to be expressed in LRFCs right before the first asymmetric cell division that hallmarks LR initiation. They continue to be expressed throughout LR primordium development, during which their expression domains become restricted to the center of the primordium. *RGI1, 4* and *5*, are believed to be the most important receptors for GLV6/10 peptides during LR initiation (Fernandez *et al*., 2020; Jourquin *et al*., 2022). *RGI1* was shown to be expressed throughout the young XPP, as well as in LRFCs and young LR primordia, while its expression is mainly restricted to the flanks of the primordia during later developmental stages. *RGI5* expression is initiated in LRFCs upon LR initiation and can still be observed after the first anticlinal cell division, but is strongly reduced in later developmental stages. These expression patterns suggest that GLV6/10 peptides can be perceived by the peptide producing LRFCs themselves, as well as by the flanking pericycle cells. Overexpression (OE) or treatment with synthetic GLV6 and GLV10 peptides inhibits LR initiation and triggers aberrant anticlinal cell divisions throughout the pericycle (Meng *et al*., 2012; Fernandez *et al*., 2015). Conversely, double *glv6glv10* knock-out mutants were found to have increased total (non-emerged + emerged) LR primordium densities and display an irregular spacing of LR primordia along the primary root, where LR primordia often occur in pairs or clusters (Fernandez *et al*., 2020; Jourquin *et al*., 2022). These findings suggest that GLV6 and GLV10 peptides serve as redundantly acting inhibitors of LR initiation that regulate the spacing of LRs along the primary root. Recently, we have characterized the GLV6/10-induced transcriptional response through an RNA-sequencing experiment designed to specifically capture the effect GLV peptide signaling during LR initiation (Jourquin *et al*., 2022). To gain a more thorough understanding of how GLV6/10 peptides regulate the spacing of LR primordia along the primary root, and given the prominent role of auxin in LR development, we set out to investigate the interplay between GLV6/10 and auxin signaling during LR initiation in more detail. Further examination of the GLV6/10-induced transcriptional response indicated that GLV6/10 peptide signaling regulates the expression of a particular subset of auxin response genes, downstream of the canonical auxin signaling pathway, and simultaneously attenuates the local accumulation of auxin in xylem-pole pericycle cells via the PIN3 and PIN7 auxin export proteins. The GLV6/10 signaling pathway thus constitutes a negative feedback loop on the formation of auxin maxima that ensures proper LR spacing.

## Materials and Methods

### Plant material and growth conditions

Unless stated otherwise, surface sterilized seeds were sown directly on solid ½ Murashige and Skoog (MS) medium (2.154g/L MS (Duchefa Biochemie), 1% sucrose, 0.1g/L Myo-inositol (Sigma-Aldrich), 0.5g/L MES (Duchefa Biochemie), 0.8% Plant Tissue Culture Agar (Lab M)). After 48h of stratification at 4°C, Petri plates were incubated at 21°C in continuous light. The following transgenic and mutant *Arabidopsis* lines were described elsewhere: *iGLV6, rgi1rgi5/iGLV6* (Fernandez *et al*., 2020), *CYCA2;4:GUS* (Vanneste *et al*., 2011), *CYCB1;1:GUS* (Ferreira *et al*., 1994), *tir1-1* (Ruegger *et al*., 1998), *afb1-3, afb2-3* (Savaldi-Goldstein *et al*., 2008), *afb3-4, tir1-1afb2-3, tir1-1afb3-4* (Parry *et al*., 2009), *arf7arf19* (Okushima *et al*., 2005), *slr-1* (Fukaki *et al*., 2002), *DR5:LUCIFERASE* (Moreno-Risueno *et al*., 2010), *iGLV6/DR5:LUCIFERASE* (Fernandez *et al*., 2020), *35S:DII-VENUS* (Vernoux *et al*., 2011), *gh3*.*123456* (Porco *et al*., 2016), *abcb1-100, abcb19-101, abcb1-100abcb19-101* (Lin and Wang, 2005), *abcb4-1* (Terasaka *et al*., 2005), *abcb21-1* (Kamimoto *et al*., 2012), *abcb1-100abcb19-101abcb21-1* (Jenness *et al*., 2019), *ABCB1:ABCB1-GFP, ABCB19:ABCB19-GFP* (Mravec *et al*., 2008), *aux1-7, AUX1:AUX1-YFP* (Swarup *et al*., 2004), *lax1, lax2, lax3* (Swarup *et al*., 2008), *pin1 (En134)* (Gälweiler *et al*., 1998), *pin2 (eir1-1)*(Roman *et al*., 1995), *pin1pin2* (Gälweiler *et al*., 1998), *pin3-2* (Friml *et al*., 2002), *pin7-1, pin3-5pin7-1* (Benková *et al*., 2003), *PIN3:PIN3-GFP, PIN7:PIN7-GFP* (Billou *et al*., 2005).

### Phenotypic analyses

To assess the sensitivity of different mutant lines to the inhibitory effect of GLV6/10 on LR initiation, Col-0 and mutant seedlings were sown on untreated medium and transferred to medium supplemented with 100nM GLV10p or untreated control medium at 3 days after germination (DAG). Synthetic GLV10p (DY(SO_3-_)PKPSTRPPRHN) was obtained from Genscript (>70% purity). Emerged LR numbers were counted on 12DAG seedlings using a stereo microscope. Total LR (non-emerged + emerged) numbers were quantified in 9DAG seedlings using an Olympus BX53 DIC microscope after clearing the roots as previously described (Malamy and Benfey, 1997). Scans of roots were used to measure primary root lengths with FIJI (Schindelin *et al*., 2012; Schneider *et al*., 2012) and densities were calculated by dividing emerged or total LR numbers by primary root lengths.

To determine whether auxin can rescue the LR inhibition caused by *GLV6*^*OE*^, *iGLV6* seedlings (containing an estradiol inducible *GLV6*^*OE*^ construct) were grown on plates with 2µM estradiol, 0.1µM 1-naphthaleneacetic acid (NAA), a combination of both, or dimethylsulfoxide (DMSO) as mock treatment. At 12DAG, emerged LR densities were quantified after which the roots were cleared as previously described (Malamy and Benfey, 1997), and imaged using an Olympus BX53 DIC microscope. The effect of GLV10p treatment on the pericycle of *arf7arf19* mutant roots, and on the primary root of *abcb1abcb19* mutants, was assessed using 9DAG seedlings grown on untreated or 100nM GLV10p-containing medium. Similarly, the effect of *GLV6*^*OE*^ on the pericycle of *slr1/iGLV6* and 10µM naphthylphthalamic acid-treated (NPA; Duchefa Biochemie) *iGLV6* roots was assessed using 9DAG seedlings grown on medium with 2µM estradiol (Sigma) or an equivalent volume of dimethylsulfoxide (DMSO). Seedlings were fixed and cleared using the ClearSee protocol (Kurihara *et al*., 2015; Ursache *et al*., 2018) and cell walls were stained using Calcofluor White (Sigma). Images were taken using a Zeiss LSM710 confocal microscope. Calcofluor White was excited at 405nm and acquired at 430-470nm.

### Expression pattern analyses

To assess the effect of GLV6/10 on *CYCA2;4* and *CYCB1;1* expression, *CYCA2;4:GUS* seedlings were grown on untreated medium or medium containing 200nM GLV10p, while *CYCB1;1* and *CYCB1;1/35S-GLV6* seedlings were all grown on untreated medium. Seedlings were collected for GUS (β-glucuronidase) staining at 9DAG. GUS staining was performed as previously described after fixation in 90% acetone at 4°C (Beeckman and Engler, 1994). Images were taken with an Olympus BX53 DIC microscope or a KEYENCE VHX-7000 digital microscope.

All confocal images were taken using a Zeiss LSM710 microscope. GFP was excited at 488nm and acquired at 495-540nm. YFP and VENUS were excited at 514nm and acquired at 515-555nm. To analyze the effect of GLV10p on fluorescent reporter lines (*35S:DII-VENUS, ABCB1:ABCB1-GFP, ABCB19:ABCB19-GFP, AUX1:AUX1-YFP, PIN3:PIN3-GFP* and *PIN7:PIN7-GFP*) during LR initiation, seedlings were gravistimulated (90° counter clockwise rotation) at 4DAG, and 8h later, transferred to untreated medium or medium with 1µM GLV10p. Root bends, older parts of the maturation zone, and root tips, were imaged 10h after peptide treatment (= 18h after gravistimulation). For measurements of PIN7-GFP intensity, *PIN7:PIN7-GFP* seedlings were transferred to untreated medium or medium with 1µM GLV10p at 8DAG. At 9DAG, confocal images were taken in the young maturation zone (from the start of the maturation zone up to the first stage II primordium) of 15 seedlings per condition. For every seedling, the mean fluorescence intensity of the PIN7-GFP signal was quantified in three anticlinal division planes within the xylem-pole pericycle using FIJI (Schindelin *et al*., 2012; Schneider *et al*., 2012).

### HPLC-MS/MS based profiling of auxin levels

Seedlings of the *iGLV6* line were grown on nylon membranes with a 20μm mesh size (Prosep) placed on top of standard ½MS medium for 4 days, after which they were gravistimulated (90° counterclockwise rotation). 8h after gravistimulation, seedlings were transferred to medium containing 2µM estradiol or an equivalent volume of DMSO. 12h and 24h after transfer, 100 root bends were dissected and collected per sample. 5 biological replicates were obtained for each condition. The extraction, purification and the LC-MS analysis of endogenous IAA and its metabolites and amino acid conjugates was carried out according to (Novák *et al*., 2012) with some modification to also include jasmonic acid + amino acid conjugates (data not shown). Briefly, 100 root bends per sample were homogenized using a bead mill (27Hz, 10min, 4°C; MixerMill, Retsch GmbH, Haan, Germany) and extracted in 1ml of 50mM sodium phosphate buffer containing 1% sodium diethyldithiocarbamate and a mixture of 13C6- or deuterium-labelled internal standards. After centrifugation (14000 RPM, 15min, 4°C), the supernatant was immediately processed as follows. Sample pH was adjusted to 2.5 using 1M HCl. Samples were applied to preconditioned Oasis HLB solid-phase extraction columns (30 mg 1 cc, Waters Inc., Milford, MA, USA). After sample application, columns were rinsed with 2ml 5% methanol. Compounds of interest were then eluted with 2ml 80% methanol. Mass spectrometry analysis and quantification were performed using an LC-MS/MS system comprised of a 1290 Infinity Binary LC System coupled to a 6495 Triple Quad LC/MS System with Jet Stream and Dual Ion Funnel technologies (Agilent Technologies, Santa Clara, CA, USA).

### DR5:LUCIFERASE imaging

To count the number of DR5:LUCIFERASE (DR5:LUC) sites per root, 9DAG *DR5:LUC* and *gh3*.*123456/DR5:LUC* seedlings were sprayed with 1 mM D-Luciferin (Duchefa Biochemie) solution [D-Luciferin dissolved in 0.01% (v/v) Tween80 and 0.1% (v/v) DMSO] and kept in total darkness for 10min before imaging. Emitted LUCEFERASE signal was captured using an ANDOR iKon-M 934 charge-coupled device (CCD) camera (Oxford Instruments) paired with a fixed lens (Spacecom 43F2409M-MP C 4/3’’ 24mm F0.9) over a 20min exposure time. Corresponding brightfield images were taken to measure the primary root lengths with FIJI (Schindelin *et al*., 2012; Schneider *et al*., 2012) and densities were calculated by dividing DR5:LUC site numbers by primary root lengths. For time-lapse imaging of *DR5*:*LUC* expression in gravistimulation-induced root bends, 4DAG *DR5:LUC/iGLV6* seedlings were gravistimulated (90° counter clockwise rotation), and 8h later, transferred to medium containing 2µM estradiol or DMSO, after which they were immediately prepared for imaging. For the comparative time lapse imaging of *DR5:LUC* expression in peptide treated wild-type and *gh3*.*123456* backgrounds, 5DAG *DR5:LUC* and *gh3*.*123456/DR5:LUC* seedlings were transferred to untreated medium or medium with 1µM GLV10p, and immediately prepared for imaging. Seedlings were sprayed with 1 mM D-Luciferin solution and kept in total darkness for 10min before initiating the time course. Emitted LUCIFERASE signal was captured using a NightSHADE LB985 in vivo plant imaging system (Berthold technologies) equipped with a deep-cooled slow scan CCD camera and the accompanying lens (Andor Instruments). An image was taken every 20min with a 10min exposure time. Measurement of signal intensities in the root bends and the generation of kymographs was performed using Fiji.

### RT-qPCR experiments

Roots were sampled from 7DAG Col-0 and *arf7arf19* seedlings that were incubated for 24h in liquid ½MS, with or without 1µM GLV10p. Total RNA was isolated using the ReliaPrep™ RNA Miniprep System (Promega). cDNA was synthesized with the qScript® cDNA SuperMix (Quantabio). RT-qPCRs were performed using SYBR® Green Mix (Roche) in a LightCycler® 480 (Roche). *CKA2* and *CDKA1* were included as housekeeping genes. The following primers were used in this study: *CK2A* (5’-GATCTTGGCGAGAGAATCGGTATC-3’; 5’-ACCACCATTAACGTGCGTCAAC-3’), *CDKA1* (5’-ATTGCGTATTGCCACTCTCATAGG-3’; 5’-TCCTGACAGGGATACCGAATGC-3’), *LBD16* (5’-TACAACGGCGGGGACAGGT-3’; 5’-GCTGCGAATCTTGCTGCTCC-3’), *PLT5* (5’-TGCCACGGCGGTTCTTTCCG-3’; 5’-TAGATCTCGGCGGCGGACGG-3’), *PUCHI* (5’-CACCACCACAACCGGCGAGA-3’; 5’-TCGGCTGCGTAACGACCCTA-3’), *AT5G09800* (5’-CGCCGTGCTTGAGTTTATCG-3’; 5’-ATGCGAGTTTCACGCGTTTT-3’), *AT3G02850* (5’-TCTCTTCGTTTCCGAGTGGC-3’; 5’-CAATCCCGCGCCTTCTTTTC-3’), *AT3G26610* (5’-ATCCAAGGCTCAGGGACTCT-3’; 5’-CCGTTGGTTTCATTGGTGGG-3’)

### Gene Ontology enrichment analyses

Overrepresented gene ontology (GO) terms for biological processes, as defined by the PANTHER classification system (Mi *et al*., 2019), were performed via a Fisher’s Exact test with Bonferroni correction using the PANTHER GO tool (www.pantherdb.org).

### Statistical analyses

Statistical analyses were performed in R (R Core Team, 2020). For LR and DR5:LUC site density data, a Poisson model (or a quasipoisson model in case of over- or under-dispersion) was fitted to the LR or DR5:LUC site counts, with treatment, genotype, and their interaction as fixed effects. A log-link function was applied, and log-transformed primary root lengths were used as an offset variable. Statistical analysis of primary root lengths was performed via a two-way analysis of variance (ANOVA) with treatment, genotype, and their interaction as fixed effects. Contrasts and post-hoc interaction analyses were set up using the “emmeans” or the “phia” package (De Rosario-Martinez, 2015; Lenth, 2021) and a Dunnett correction was implemented where applicable. Fluorescence intensity measurements were compared via a Mann-Whitney test. For the statistical analysis of RT-qPCR data, log2 fold changes in expression levels between treatment and mock conditions were calculated for each genotype, and two-tailed Student’s t-tests were performed to compare treatment effects between genotypes. The levels of free IAA and its derivatives were compared via two-way ANOVA with treatment and time as fixed effects, followed by post-hoc Tukey tests. For the pairwise comparison of transcriptomic datasets, the probability of finding the observed number of overlapping genes (within the total number of genes for which reads/hits were found in all experiments) was determined using a hypergeometric test.

## Results

### GLV6/10 signaling regulates a subset of auxin response genes downstream of the canonical auxin signaling pathway

We previously identified a set of about 800 genes that are differentially regulated upon *GLV6* overexpression (OE) during LR initiation via an RNA-sequencing experiment using an estradiol-inducible *GLV6*^OE^ line (*iGLV6*) in combination with gravistimulation-induced LR initiation (Jourquin *et al*., 2022). In agreement with a regulatory role for GLV6/10 peptides during LR initiation, GO-enrichment analysis on these GLV response genes revealed an enrichment for genes associated with LR formation and root system architecture, as well as auxin responses and transport (Supplementary Fig. 1S). Indeed, the previously established target genes of the GLV6/10-RGI module, *PUCHI, PLT5* and *LBD16*, are all known to be auxin inducible (Hirota *et al*., 2007; Okushima *et al*., 2007; Hofhuis *et al*., 2013). Comparative analysis of the GLV6-induced transcriptional response with previously reported transcriptomic effects of NAA and IAA treatment on *Arabidopsis* roots (Okushima *et al*., 2005; Vanneste *et al*., 2005) revealed that nearly all GLV6-regulated genes are also regulated by auxin in a SLR/IAA14 and ARF7/19 dependent manner (Fig. 1A). More precisely, most GLV6-induced genes are also induced by auxin, while the large majority of genes downregulated by GLV6 are downregulated by auxin as well (Supplementary Fig. S2, Supplementary Table 1).

**Fig. 1.**
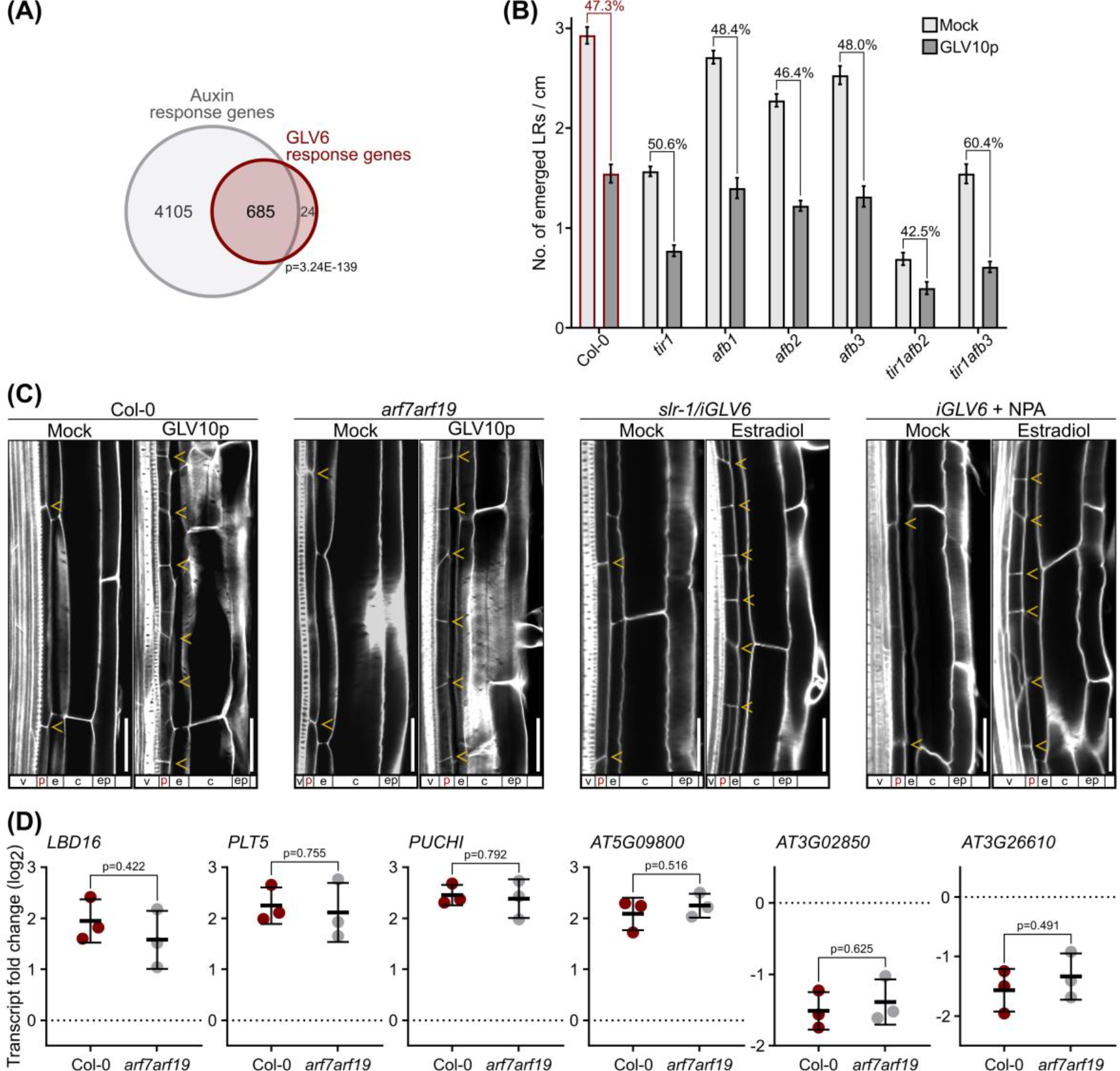
GLV6/10 peptide signaling induces a partial auxin response downstream of the canonical auxin signaling pathway. **(A)** Venn diagram showing the overlap between GLV6-regulated genes and the genes found to respond to NAA and/or IAA treatment in roots of Col-0, *slr-1* and *arf7arf19* seedlings as shown by the microarray experiments performed by Vanneste *et al*.,*2005* and Okushima *et al*., *2005*. A list of these genes and their transcript fold changes can be found in *supplementary table 1*. Statistical significance was determined via a hypergeometric test. **(B)** Emerged LR densities of 12-day old wild-type, *tir1, afb1, afb2, afb3, tir1afb2 and tir1afb3* mutant seedlings, treated or not with 100nM GLV10p. The mean numbers of emerged LRs per centimeter ± SE are shown. For each genotype, the percentual decrease in LR density as a result of GLV10p treatment is indicated. Asterisks indicate a decrease that is significantly different (p-value < 0.05) from that observed in the wild-type control (none in this case), n = 15. Statistical significance was determined via quasipoisson regression with Dunnett’s correction. **(C)** Confocal images of longitudinal sections through the maturation zone of 9-day old Col-0, *arf7arf19, slr-1/iGLV6* and NPA-treated (10µM) *iGLV6* roots, treated or not with 100nM GLV10p or 2 µM estradiol (to induce *GLV6* overexpression) as indicated above each image. Anticlinal divisions within a single pericycle cell file are indicated with yellow arrowheads. The different tissue layers are indicated below each image; v=vasculature, p=pericycle, e=endodermis, c=cortex and ep=epidermis. Cell walls were stained with Calcofluor White. Bars, 20µm. **(D)** Log2 fold changes (FCs) in expression levels of *GLV6* response genes in wild-type and *arf7arf19* mutant roots upon treatment with 1µM GLV10p. Individual datapoints and mean ± SD are shown, n = 3. Dotted lines indicate log2(FC) = 0 (i.e. no change in expression level). Statistical significance was determined using Student’s t-tests.

This large overlap between GLV6 and auxin response genes, prompted us to investigate whether GLV6/10 peptides establish their effects via auxin. When grown in the presence of GLV10p, mutants in the auxin receptor TRANSPORT INHIBITOR RESPONSE 1 (TIR1), or its homologous AUXIN SIGNALING F-BOX PROTEINS (AFBs) (Parry *et al*., 2009), showed a similar percentual decrease in emerged LR density compared to wild-type seedlings (Fig. 1B), suggesting that the perception of auxin is not necessary for the inhibitory effect of GLVs on LR initiation. Auxin largely regulates LR development via the ARF7 and ARF19 transcription factors, and their negative regulator SLR/IAA14 (Fukaki *et al*., 2002; Okushima *et al*., 2005). Since *arf7arf19*, as well as the dominant negative *slr-1* mutants, do not produce LR primordia, the effect of GLV10p treatment or *GLV6*^*OE*^ on these mutants was assessed through microscopic analysis of the pericycle. This revealed that both mutants still show the ectopic anticlinal pericycle cell divisions typically observed in the presence of an excess of GLV6/10 peptides (Fig. 1C). Furthermore, such ectopic anticlinal pericycle divisions can also still be induced upon *GLV6*^*OE*^ in the presence of the auxin transport inhibitor NPA (Fig. 1C), which typically inhibits LR initiation by preventing auxin accumulation in LRFCs (Casimiro, 2001). These data thus indicate that the effect of GLV peptides on pericycle cell divisions does not require auxin transport or signaling. In agreement with this, RT-qPCR analysis of a number of *GLV6* response genes indicated that their response to GLV10p treatment is not attenuated in the *arf7arf19* mutant background (Fig. 1D). Combined with the fact that expression of *GLV6* and *GLV10* is known to be auxin inducible via ARF7/19 and their downstream target LBD16 (Fernandez *et al*., 2020; Jourquin *et al*., 2022), this suggests that these GLV peptides are involved in the regulation of a part of the transcriptional auxin response, downstream of the core auxin signaling machinery that controls LR initiation.

The overlapping GLV- and auxin-induced transcriptional responses seem somewhat counterintuitive, since auxin treatments typically trigger formative divisions throughout the xylem-pole pericycle, resulting in the formation of many LRs along the whole primary root (Parizot *et al*., 2008), while GLV6/10 peptides inhibit LR initiation. However, an excess of GLV6/10 peptides does not merely inhibit the first asymmetric cell divisions of LRFCs. Rather, as evidenced by the long stretches of short pericycle cells observed in peptide-treated roots, an overabundance of GLV6/10 peptides triggers ectopic anticlinal cell divisions throughout the pericycle (Fig. 1C)(Meng *et al*., 2012; Fernandez *et al*., 2015, 2020). In agreement with this, the late transcriptional response to *GLV6*^OE^ (6h after induction) is enriched for genes involved in the regulation of cell division (Supplementary Fig. S3). The cell-cycle gene that was most strongly induced after *GLV6*^*OE*^ was *CYCA2;4*, and analysis of a *CYCA2;4:GUS* reporter line revealed that it is ectopically expressed along the whole pericycle after treatment with GLV10p, as opposed to its typical association with sites of LR development in untreated conditions (Fig. 2A, 2B) (Vanneste *et al*., 2011). In contrast, the cell cycle gene *CYCB1;1*, which typically marks the formative pericycle cell divisions of LR founder cells and LR primordia, and which is known to be induced throughout the pericycle upon auxin treatment (Himanen *et al*., 2002), did not respond to *GLV6*^*OE*^ in our RNA-sequencing experiment, nor could any *CYCB1;1* promoter driven GUS activity be observed in the pericycle of *GLV6* overexpressing plants (Fig. 2C). Despite the overlapping transcriptional responses, there thus seems to be an important difference between the pericycle divisions triggered by auxin, which are formative and lead to the development of LR primordia, and the divisions triggered by an excess of GLV6/10 peptides, which do not give rise to LR primordia. In this context, it should be noted that the GLV6 response genes constitute only a fraction of the complete auxin response that was characterized through prior transcriptome analyses in the root (Fig. 1A), indicating that *GLV6*^*OE*^ only triggers a partial auxin response. This GLV-induced partial auxin response thus seems to stimulate pericycle cell divisions, but is not sufficient for the proper initiation of LR development.

**Fig. 2.**
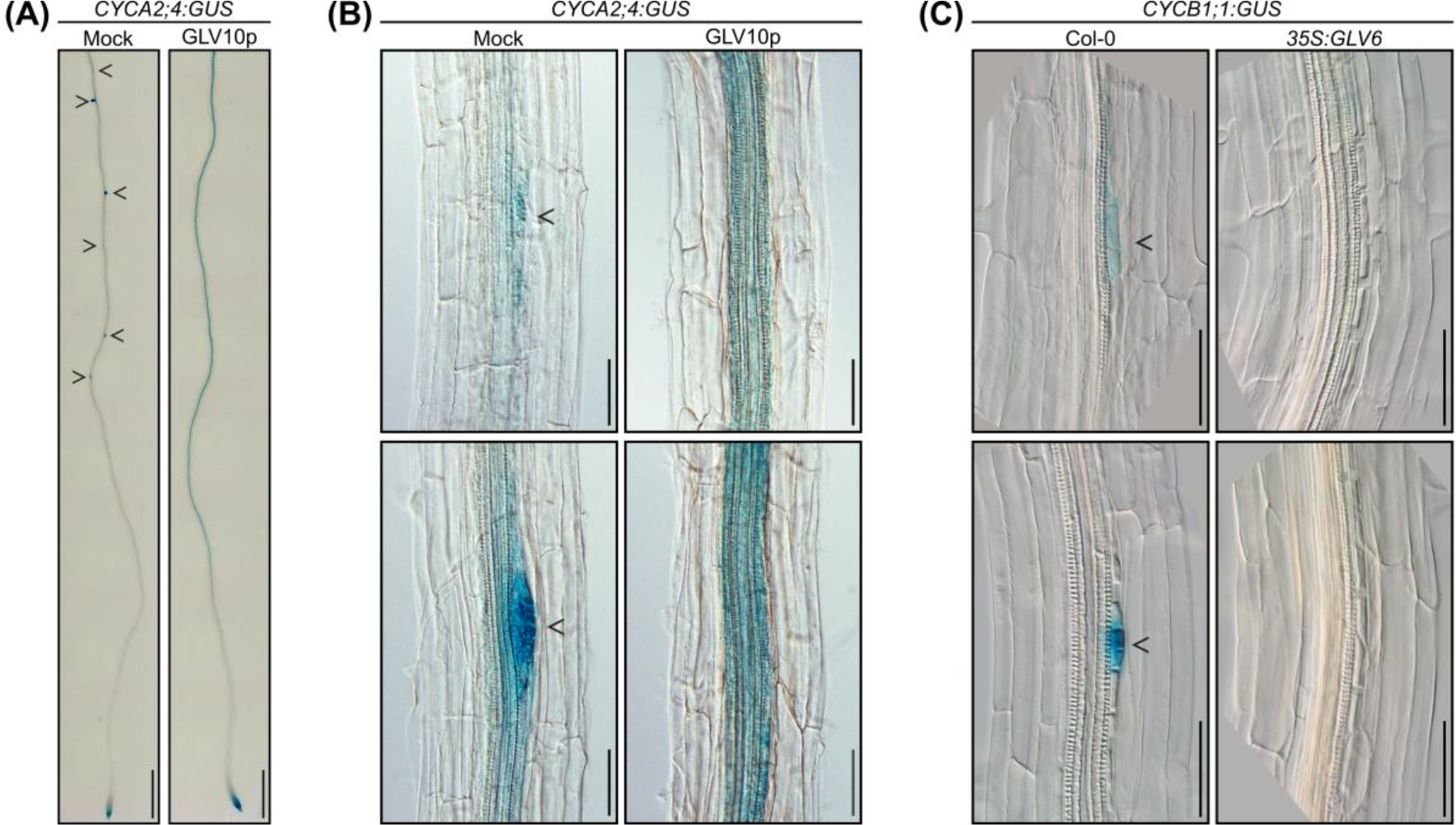
The GLV6/10-induced pericycle divisions differ from those induced by auxin during LR initiation. **(A)** *CYCA2;4:GUS* expression pattern in *Arabidopsis* roots treated or not with 200nM GLV10p. LR primordia are highlighted with black arrowheads. Scale bars represent 0.25cm. **(B)** Close-ups of *CYCA2;4:GUS* expression in *Arabidopsis* roots treated or not with 200nM GLV10p. LR primordia are highlighted with black arrowheads. Scale bars represent 50µm. **(C)** Close-ups of *CYCB1;1:GUS* expression in wild-type and *35S:GLV6 Arabidopsis* roots. LR primordia are highlighted with black arrowheads. Scale bars represent 50µm.

### GLV6/10 overabundance disrupts the auxin accumulation in LRFCs required for lateral root initiation

Given the opposing effects of GLV6/10 and auxin on LR initiation, despite their overlapping transcriptional effects, we investigated the effect of GLV6/10 signaling on auxin signaling and homeostasis. Our previous analysis of the expression of the auxin-responsive *DR5* promoter suggested that *GLV6*^*OE*^ attenuates auxin signaling activity in pre-branch sites (Fernandez *et al*., 2020). A more thorough analysis indeed revealed that, when *GLV6*^*OE*^ is induced 8 hours after gravistimulation, the DR5:LUCIFERASE (DR5:LUC) signal in the gravistimulation-induced pre-branch site is gradually reduced, while a clear pre-branch site with high DR5:LUC intensity is typically established and maintained in the root bends of untreated seedlings (Fig. 3A, 3B). *GLV6*^*OE*^ thus seems to abolish pre-branch site formation completely. Additionally, expression of the *35S:DII-VENUS* reporter was analyzed upon GLV10p treatment after gravistimulation-induced LR initiation. The fusion of VENUS to the degron domain of an AUX/IAA protein (DII) renders it vulnerable to auxin-induced degradation via interaction with TIR1/AFB proteins, making its fluorescence intensity inversely correlated to endogenous auxin levels (Vernoux *et al*., 2011). In the absence of synthetic GLV10p, only a very faint DII-VENUS signal could be observed in the pericycle after gravistimulation-induced LR initiation, while a clear nuclear DII-VENUS signal was detected in pericycle cells of GLV10p-treated seedlings (Fig. 3C). Hence, these data imply that *GLV6*^*OE*^ and GLV10p treatments disrupt the accumulation of auxin in LRFCs, thereby preventing LR initiation. In agreement with this, treatment with the synthetic auxin analog NAA is able to partially restore LR development when *GLV6* is being overexpressed. Emerged LR densities were quantified upon estradiol-induced *GLV6*^*OE*^ in combination with NAA treatment (Fig. 3D). The high LR densities typically observed in the presence of NAA were clearly reduced upon *GLV6*^*OE*^, but were still high compared to the nearly complete inhibition of LR development observed when *GLV6* is overexpressed in the absence of NAA. Importantly, microscopic analysis of these NAA-treated roots showed that the pericycle cells between LR primordia still undergo many ectopic cell divisions when *GLV6* is being overexpressed, while the LR primordia in roots treated with NAA alone are flanked by pericycle cells that are long and narrow as observed in untreated conditions (Fig. 3E). This suggests that NAA treatment does not inhibit the occurrence of GLV6/10-induced ectopic pericycle cell divisions, but rather overcomes or overrules the inhibitory effect on LR initiation, possibly by compensating for the GLV6/10-induced lack of auxin in LRFCs.

**Fig. 3.**
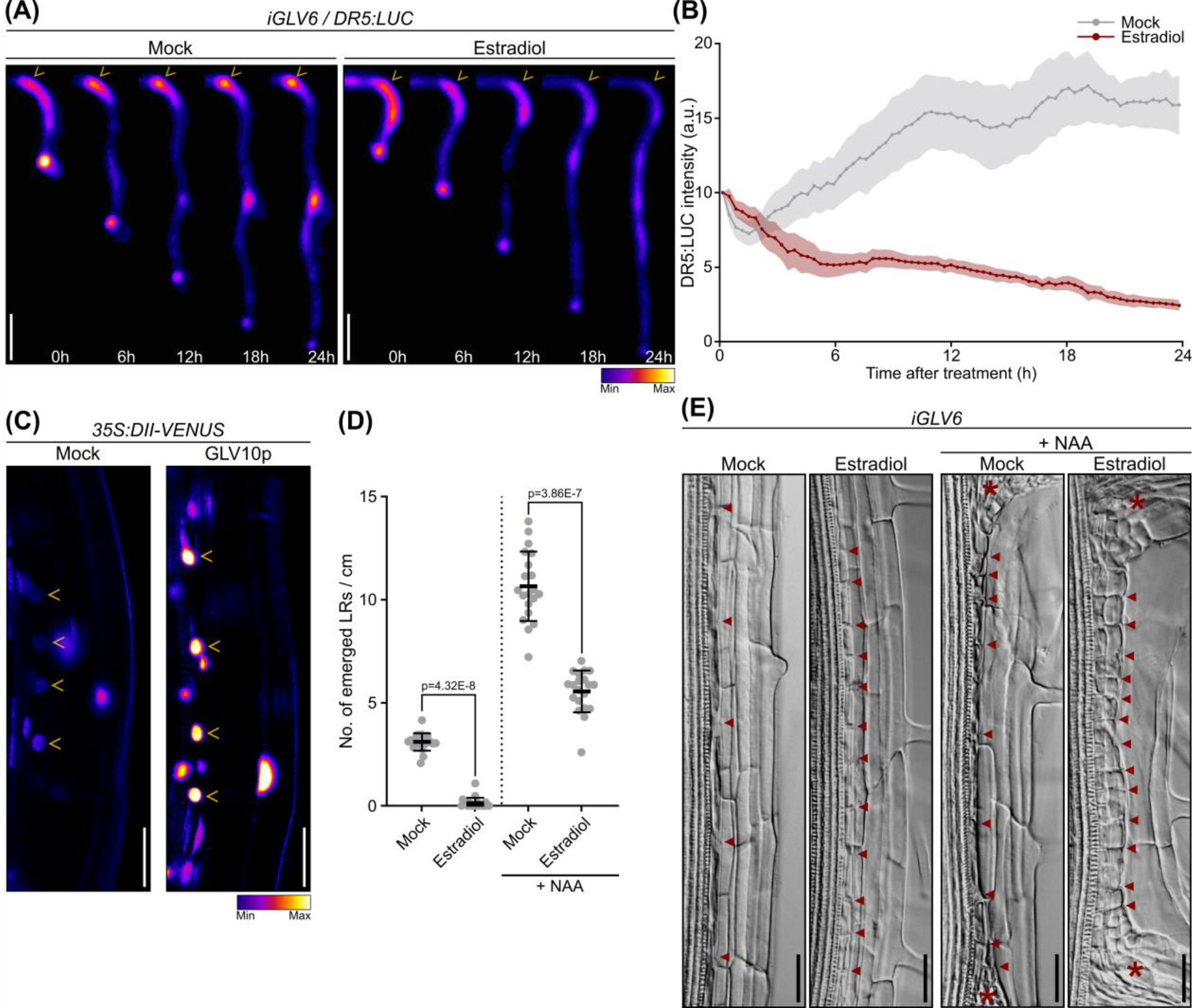
GLV6/10 overabundance disrupts the accumulation of auxin in LRFCs required for LR initiation. **(A)** Snapshots of a 24h time course capturing LUCIFERASE activity in untreated and estradiol-treated *iGLV6/DR5:LUCIFERASE* roots. Roots were gravistimulated 8h prior to the start of the time series and *GLV6* overexpression was instigated at the onset of the time course. The locations of pre-branch sites in the root bend are marked with yellow arrowheads. LUCIFERASE intensity is shown in arbitrary units. Scale bars represent 0.2cm. **(B)** Time course of LUCIFERASE intensity in pre-branch sites in gravistimulation-induced root bends of *iGLV6/DR5:LUC* roots treated or not with estradiol. Roots were gravistimulated 8h prior to the start of the time series and *GLV6* overexpression was instigated at the onset of the time course. LUCIFERASE intensity in each gravistimulation-induced pre-branch site was measured every 20min for 24h, and normalized against its intensity at the start of the time series, which was arbitrarily placed at 10 arbitrary units (a.u.). The mean ± SE are shown, n = 10. **(C)** Representative confocal images of *35S:DII-VENUS* expression in gravistimulation-induced root bends. Roots were treated or not with 1µM GLV10p at 8h after gravistimulation and imaged 18h after gravistimulation. Nuclei of pericycle cells are indicated with white arrowheads. Scale bars represent 30µm. **(D)** Emerged LR densities of 12-day old *iGLV6* seedlings treated or not with 0.1µM NAA and/or 2µM estradiol to induce *GLV6* overexpression. Individual datapoints and mean ± SD are shown, n = 20. Statistical significances were determined via quasipoisson regressions. **(E)** DIC images of longitudinal sections through the maturation zone of 12-day old *iGLV6* roots treated or not with 0.1µM NAA and/or 2µM estradiol to induce *GLV6* overexpression. Anticlinal divisions within a single pericycle cell file are indicated with red arrowheads. LR primordia are indicated with a red asterisk. Scale bars represent 30µm.

### Exploring the effect of GLV6/10 peptide signaling on auxin conjugation during lateral root initiation

The set of GLV6 response genes, identified through the *GLV6*^*OE*^ RNA-sequencing experiment, contains several genes involved in auxin metabolism and transport that might be responsible for the disruption of auxin accumulation in LRFCs when GLV6/10 peptides are in overabundance. This includes four GH3 encoding genes (*GH3*.*1, GH3*.*2, GH3*.*3* and *GH3*.*5*), which were all found to be downregulated upon *GLV6*^*OE*^ in an RGI-dependent manner (Supplementary Fig. S4). In fact, they are among the few genes that show opposing responses to *GLV6*^*OE*^ and auxin treatment (Supplementary Table 1), since they are typically induced by auxin rather than suppressed (Hagen and Guilfoyle, 2002; Okushima *et al*., 2005). As members of the group II GH3s, they function as enzymes that catalyze the conjugation of free IAA to amino acids (AAs) (Staswick *et al*., 2002, 2005). These IAA-AA conjugates are believed to serve as inactive storage forms of auxin and/or intermediates in the auxin degradation pathway (Ludwig-Müller, 2011; Hayashi *et al*., 2021; Casanova-Sáez *et al*., 2022). To assess the importance of GH3s for the effect of GLV6/10 on the accumulation of auxin in LRFCs, we made use of a *gh3*.*123456* sextuple mutant (Porco *et al*., 2016). In agreement with the auxin conjugating activity of GH3.1-6, the *gh3*.*123456* mutant was previously shown to contain increased levels of free auxin and displayed a higher emerged LR density compared to wild-type seedlings (Porco *et al*., 2016). We analyzed DR5:LUC expression in wild-type and *gh3*.*123456* mutant backgrounds when grown in the presence of GLV10p. As with *GLV6*^*OE*^, high concentrations of GLV10p largely inhibited the establishment of pre-branch sites and could even reduce the DR5:LUC activity in pre-branch sites that were established before treatment (Fig. 4A). In *gh3*.*123456* mutants on the other hand, pre-branch sites with clear DR5:LUC expression could still be established after GLV10p treatment (Fig. 4A). In agreement with this, *gh3*.*123456* mutant seedlings grown in the presence of GLV10p showed a significantly less pronounced reduction in DR5:LUC site density compared to wild-type seedlings (Fig. 4B). Furthermore, the GLV10p-triggered reductions in emerged, as well as total LR densities, were significantly less severe in *gh3*.*123456* mutants than in wild-type plants (Fig. 4C, 4D). This reduced responsivity of *gh3*.*123456* mutants to GLV10p treatment suggests that the inhibitory effect of GLV peptides on LR initiation is partially dependent on the activity of GH3 enzymes. If this would indeed be the case, GLV6/10 peptide signaling might regulate free auxin levels through GH3-mediated conjugation of auxin to AAs. To investigate this possibility, the effect of GLV peptide signaling on free IAA and IAA-AA conjugate levels was determined after gravistimulation-induced LR initiation. To this end, *iGLV6* seedlings were treated with estradiol 8 hours after gravistimulation, and root bends were sampled 12h and 24h after treatment, time points at which the DR5:LUC signal in the root bends was shown to be clearly reduced or had disappeared altogether (Fig. 3A, 3B). The HPLC-MS/MS based profiling of free IAA and its derivatives allowed for the detection of IAA itself, its inactive breakdown product oxIAA, both of their conjugates to glucose, and to the two AA conjugates, IAA-Aspartate and IAA-Glutamate. Other IAA-AA conjugates could not be detected, but as IAA-Aspartate and IAA-Glutamate are the main IAA-AA conjugates that are produced by GH3s *in planta*, changes in their levels can be expected if the activity or abundance of GH3 enzymes would be modulated (Staswick *et al*., 2005; Park *et al*., 2007; Mellor *et al*., 2016; Porco *et al*., 2016). However, *GLV6*^*OE*^ did not induce any significant changes in free IAA levels, nor in the levels of any of its derivatives or conjugates, compared to the levels in the corresponding mock treated root bends (Fig. 4E). Free auxin levels within the sampled root bends thus appear to be unaffected when *GLV6* is overexpressed. The GLV6/10-induced reduction in DR5:LUC activity in pre-branch sites is therefore likely caused by a failure to concentrate the available auxin in LRFCs, rather than by an effect on free auxin levels via GH3s. The diminished responsivity of *gh3*.*123456* mutants to GLV10p treatments might thus be caused by an indirect effect of the lack of IAA-AA conjugation on auxin transport and/or the elevated free IAA levels in *gh3*.*123456* mutant seedlings compared to wild-type plants (Porco *et al*., 2016).

**Fig. 4.**
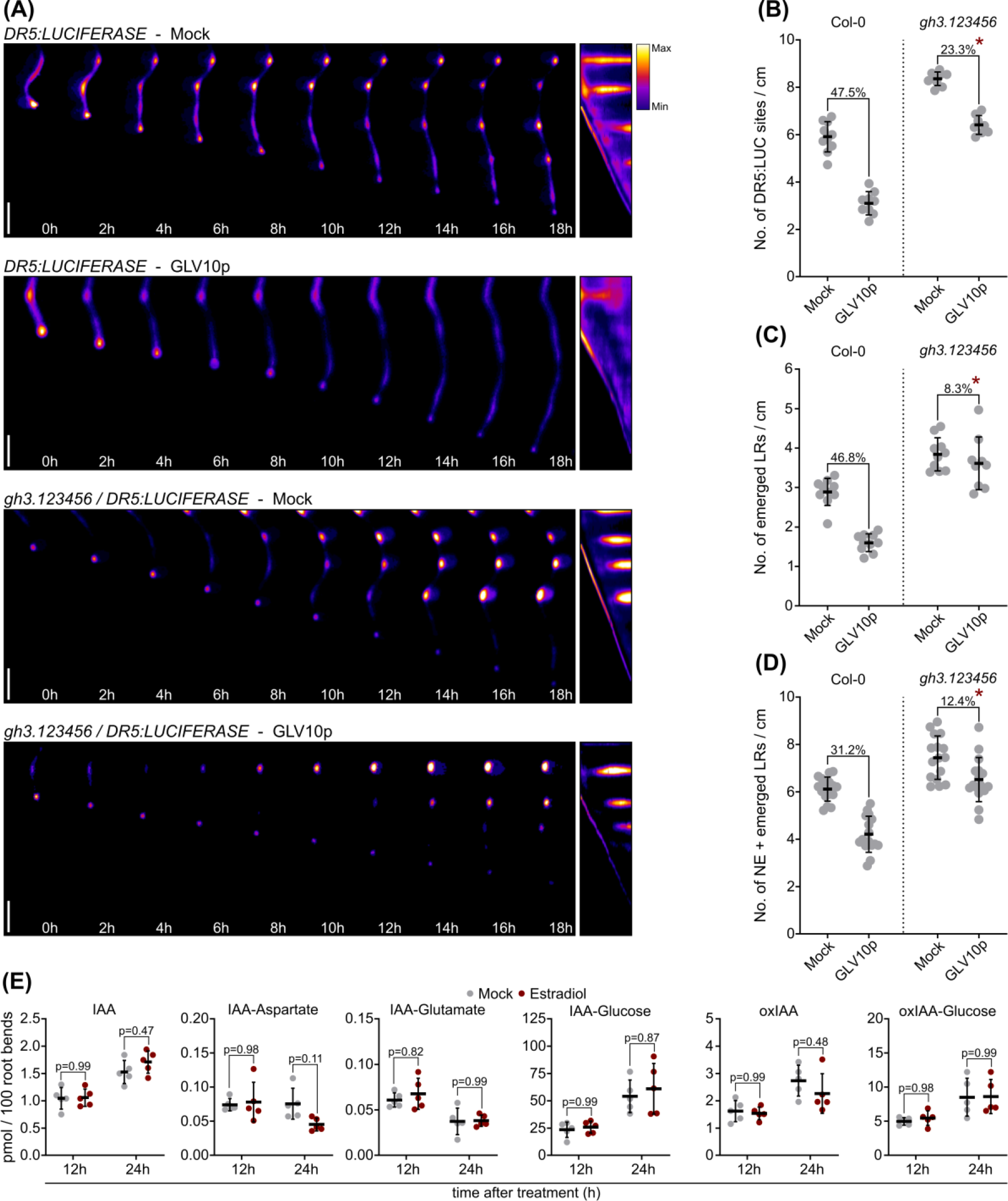
Exploring the effect of GLV6/10 peptide signaling on auxin conjugation. **(A)** Snapshots of an 18h time course capturing LUCIFERASE activity in *DR5:LUCIFERASE* and *gh3*.*123456/DR5:LUCIFERASE* roots, treated or not with 1µM GLV10p at the onset of the experiment. Panels at the far right show kymopgraphs corresponding to the depicted roots. LUCIFERASE intensity is shown in arbitrary units. Scale bars represent 0.2cm. **(B)** DR5:LUCIFERASE site densities in roots of 9-day old wild-type and *gh3*.*123456* mutant seedlings, treated or not with 100nM GLV10p. Individual datapoints and mean ± SD are shown. For each genotype, the percentual decrease in DR5:LUCIFERASE site density as a result of GLV10p treatment is indicated. Asterisks indicate a decrease that is significantly different from that observed in the wild-type control (p-value < 0.05), n = 10. Statistical significance was determined via quasipoisson regression. **(C)** Emerged LR densities of 12-day old wild-type and *gh3*.*123456* mutant seedlings, treated or not with 100nM GLV10p. Individual datapoints and mean ± SD are shown. For each genotype, the percentual decrease in emerged LR density as a result of GLV10p treatment is indicated. Asterisks indicate a decrease that is significantly different from that observed in the wild-type control (p-value < 0.05), n = 10. Statistical significance was determined via quasipoisson regression. **(D)** Total (non-emerged (NE) + emerged) LR primordium densities of 12-day old wild-type and *gh3*.*123456* mutant seedlings, treated or not with 100nM GLV10p. Individual datapoints and mean ± SD are shown. For each genotype, the percentual decrease in total LR density as a result of GLV10p treatment is indicated. Asterisks indicate a decrease that is significantly different from that observed in the wild-type control (p-value < 0.05), n=15. Statistical significance was determined via quasipoisson regression. **(E)** LC-MS/MS based profiling of the amount of IAA (indole-3-acetic acid) and its detectable derivatives in 100 root bends of gravistimulated *iGLV6* seedlings after 12h and 24h of mock (DMSO) or estradiol (2µM) treatment to induce *GLV6* overexpression. oxIAA = 2-oxindole-3-acetic acid, n = 5. Statistical significances were assessed via a two-way ANOVA and a post-hoc Tukey test.

### GLV6/10 signaling affects auxin transport during lateral root initiation via PIN auxin exporters

The inhibitory effect of GLV6/10 peptide signaling on the establishment of an auxin maximum in LRFCs might thus result from a disruption of auxin transport rather than from changes in auxin levels. The genes encoding the auxin exporters ATP-BINDING CASETTE TYPE-B 19 (ABCB19) and ABCB21 (Geisler *et al*., 2005; Kamimoto *et al*., 2012) were found to be upregulated upon *GLV6*^*OE*^ (Supplementary Fig. S4), suggesting that they might be involved in the effect of GLV6/10 peptides on auxin distribution. To assess whether these ABCBs indeed play a role in GLV6/10 peptide signaling during LR initiation, the effect of GLV10p treatment on root architecture was determined in single and higher order mutants of *ABCB19, ABCB21*, and their closest homologs *ABCB1* and *ABCB4*, respectively. Most mutants showed a GLV10p-induced decrease in emerged LR density comparable to wild-type seedlings (Fig. 5A), as well as a slight increase in primary root length typically observed upon GLV10p treatment or *GLV6*^*OE*^ (Fig. 5B). However, *abcb1abcb19* and *abcb1abcb19abcb21* mutants showed a striking decrease in primary root length upon GLV10p treatment (Fig. 5B, 5C), to such an extent that a reliable analysis of LR densities was not possible. Microscopic analysis revealed that GLV10p treatment on *abcb1abcb19* mutants caused extremely disorganized divisions, not only in the pericycle, but also in endodermal, cortical and epidermal cells, and possibly also in the vasculature (Fig. 5D). In untreated *abcb1abcb19* mutants, some aberrant divisions can be observed in the root apical meristem, but these do not seem to have a strong effect on root growth and don’t lead to any obvious defects in the maturation zone. Conversely, the GLV10p-triggered disorganization of cell divisions could only be observed in the maturation zone, while an exacerbation of the defects in the root apical meristem was not observed.

**Fig. 5.**
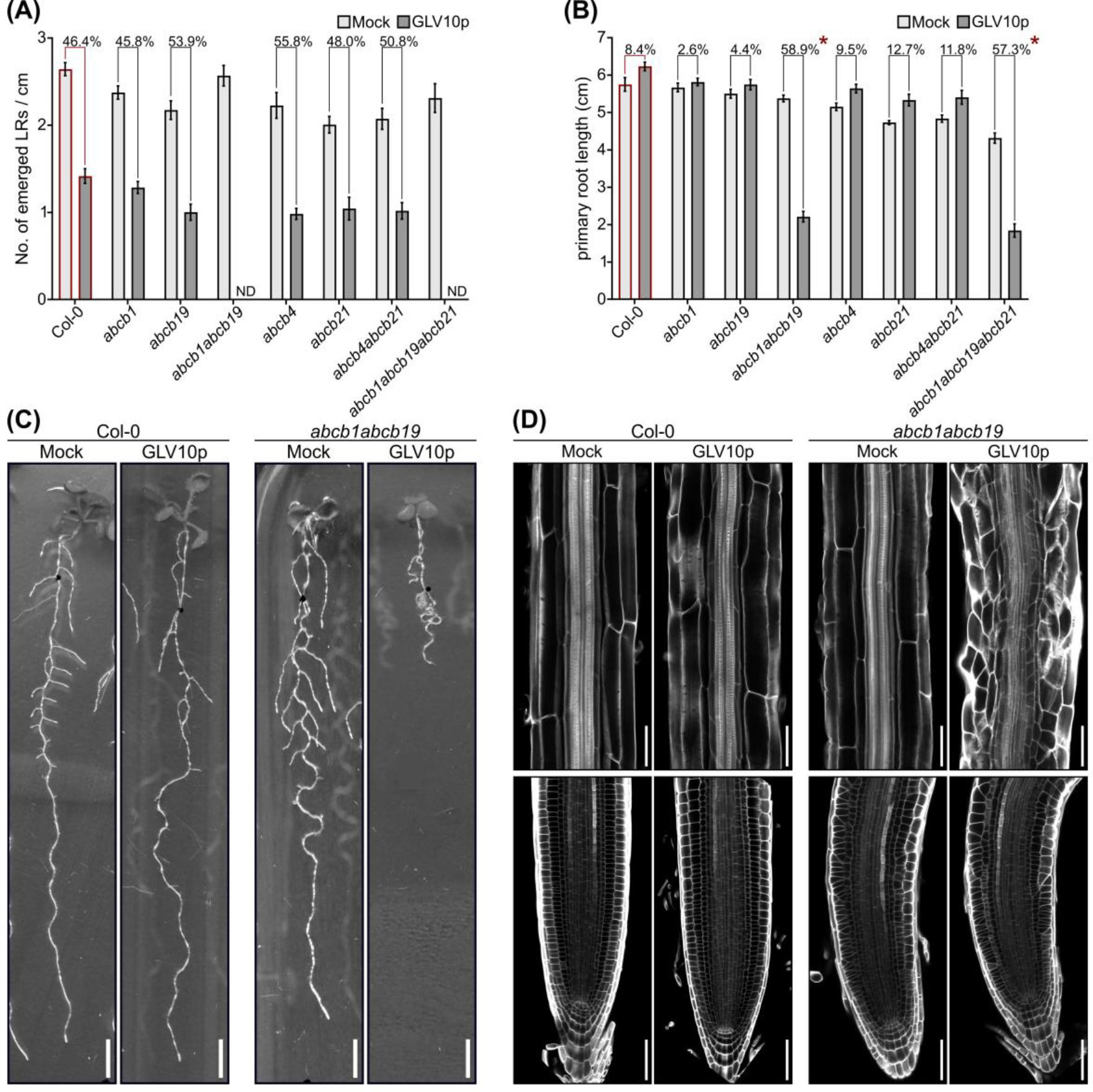
An excess of GLV peptides results in severely stunted primary root growth in *abcb1abcb19* auxin transport mutants. **(A)** Emerged LR densities of 12-day old wild-type and several *abcb* mutant lines, treated or not with 100nM GLV10p. The mean numbers of emerged LRs per centimeter ± SE are shown. For each genotype, the percentual decrease in LR density as a result of GLV10p treatment is indicated. Asterisks indicate a decrease that is significantly different (p-value < 0.05) from that observed in the wild-type control (none in this case), n = 15. Statistical significance was determined via quasipoisson regression with Dunnett’s correction. **(B)** Primary root lengths of 12-day old wild-type and several *abcb* mutant lines, treated or not with 100nM GLV10p. Mean root length ± SE are shown. For each genotype, the percentual in- or decrease in primary root length as a result of GLV10p treatment is indicated. Asterisks indicate an in- or decrease that is significantly different from that observed in the wild-type control (p-value < 0.05). Statistical significance was determined via a two-way ANOVA and a post-hoc Dunnett’s test, n=15. **(C)** Representative images of the root system of 12-day old wild-type and *abcb1abcb19* mutant seedlings, treated or not with 100nM GLV10p. Scale bars represent 0.5cm. **(D)** Confocal images of longitudinal sections through the maturation zone (upper panels) and the root tip (lower panels) of 9-day old wild-type and *abcb1abcb19* mutant roots, treated or not with 100nM GLV10p. Cell walls were stained with Calcofluor White. Scale bars represent 30µm.

The extreme response of seedlings that are defective in both *ABCB1* and *ABCB19* indicates that the combined effect of these mutations and excessive GLV peptide signaling on auxin transport results in a strong disruption of root development. However, treatment with GLV10p did not cause any obvious changes in the ABCB1-GFP and ABCB19-GFP expression patterns (Supplementary Fig. S5). It is thus currently not clear whether GLV6/10 signaling affects auxin transport through these genes.

To further examine the link between GLV6/10 peptide signaling and auxin transport, the effect of GLV10p treatment on LR development was analyzed in several other mutants for known auxin transporters. None of these mutants displayed the severe primary root growth defects observed in peptide-treated *abcb1abcb19* mutants (Supplementary Fig. S6). However, mutants in the auxin importer AUXIN RESISTANT 1 (AUX1) (Marchant *et al*., 2002; Yang *et al*., 2006) showed a stronger GLV10p-induced decrease in emerged LR density compared to wild-type seedlings, suggesting that these mutants are more responsive to the peptide treatment (Fig. 6A). Conversely, double mutants in the closely related PIN-FORMED 3 (PIN3) and PIN7 auxin exporters were found to be almost completely resistant to the inhibitory effect of GLV10p treatment on LR initiation, suggesting that GLV6/10 peptides may exert their effect on auxin transport via these PIN proteins (Fig. 6A). The expression patterns of some of these transporters also seemed to be effected by GLV10p treatment (Fig. 6B). Throughout the maturation zone of the primary root, weak *AUX1:AUX1-YFP* expression could be detected in the vasculature, but a clear signal was detected in LRFCs and developing LR primordia (Marchant *et al*., 2002; Laskowski *et al*., 2008). In the presence of GLV10p however, AUX1-YFP signal was found throughout large parts of the pericycle, in which ectopic anticlinal divisions could often be observed (Fig. 6B). Strong expression of *PIN3:PIN3-GFP* and *PIN7:PIN7-GFP* reporters was observed in the vasculature, and clear but weaker PIN3-GFP and PIN7-GFP signals were also consistently detected throughout the pericycle, which became stronger after LR initiation (Billou *et al*., 2005). An obvious and consistent effect of GLV10p treatment on *PIN3:PIN3-GFP* expression could not be discerned, but a brighter PIN7-GFP signal was observed along large parts of the pericycle, again often in association with GLV10p-induced ectopic anticlinal cell divisions (Fig. 6B). To corroborate this, the effect of GLV10p treatment on *PIN7:PIN7-GFP* fluorescence intensity was quantified in anticlinal membranes of xylem-pole pericycle cells within the young maturation zone (Fig. 6C, 6D). The significant increase in fluorescence intensity upon GLV10p treatment indicates that GLV6/10 signaling enhances PIN7 abundance in the plasma membrane of xylem-pole pericycle cells. The effect of GLV6/10-RGI signaling on auxin fluxes during LR initiation is therefore likely established via the modulation of PIN7 levels in XPP and LRFCs.

**Fig. 6.**
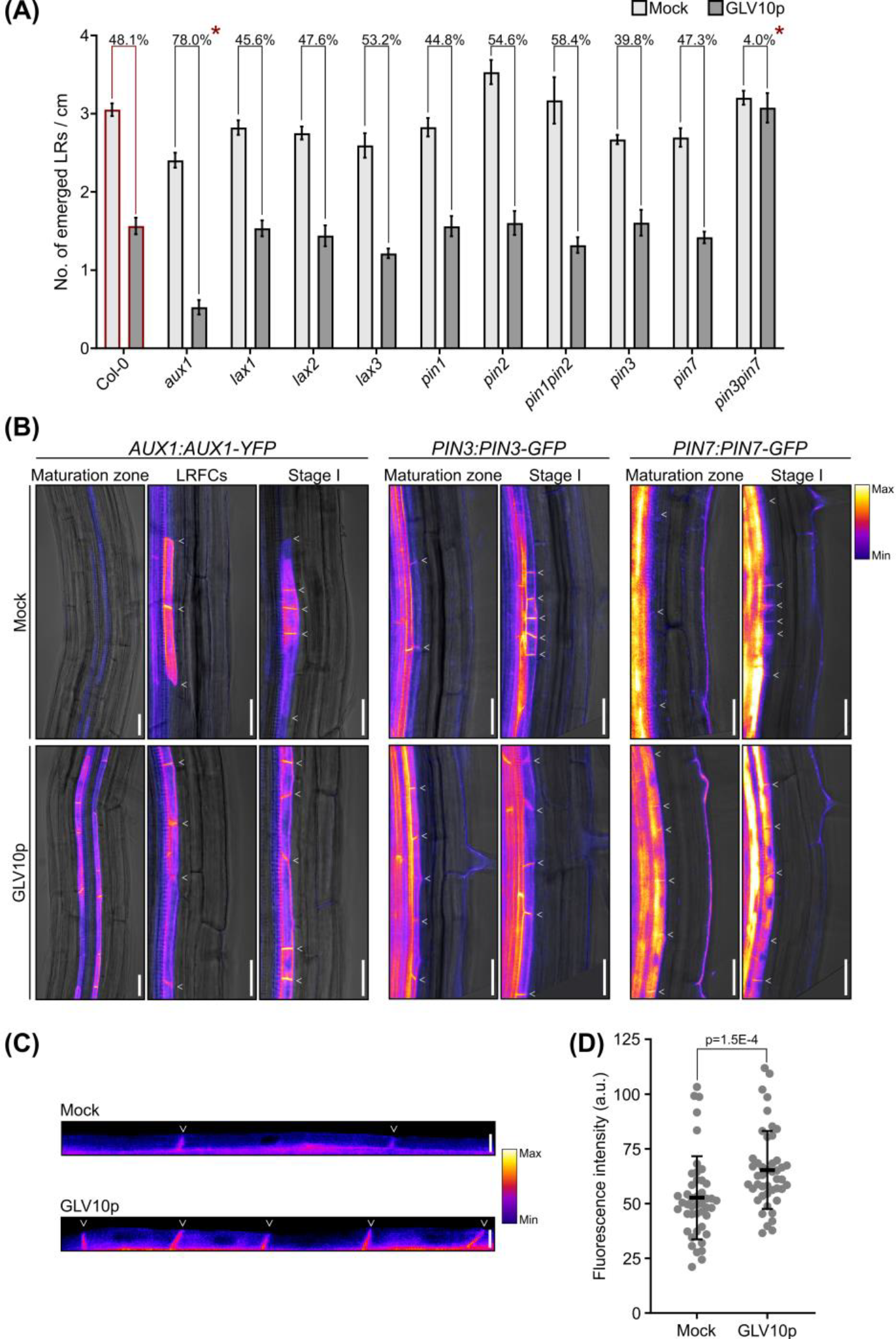
Investigating the importance of AUX1(-like) and PIN auxin transporters for GLV6/10 signaling during LR initiation. **(A)** Emerged LR densities of 12-day old wild-type and several *aux1(-like)* and *pin* mutant lines, treated or not with 100nM GLV10p. The mean numbers of emerged LRs per centimeter ± SE are shown. For each genotype, the percentual decrease in LR density as a result of GLV10p treatment is indicated. Asterisks indicate a decrease that is significantly different (p-value < 0.05) from that observed in the wild-type control, n = 15. Statistical significance was determined via quasipoisson regression with Dunnett’s correction. **(B)** Confocal images of *AUX1:AUX1-YFP, PIN3:PIN3-GFP* and *PIN7:PIN7-GFP* expression in gravistimulated roots that were treated or not with 1µM GLV10p at 8h after gravistimulation and imaged 18h after gravistimulation. Images were taken from sections of the maturation zone above the root bends and sections within the root bends. Anticlinal cell divisions within a single pericycle cell file (undergoing LR initiation or GLV10-induced aberrant divisions) are indicated with white arrowheads. Scale bars represent 30µm. **(C)** Confocal images of a small section of the xylem-pole pericycle in the young maturation zone of 9-day old *PIN7:PIN7-GFP* seedlings, grown in the absence or presence of 1µM GLV10p. White arrowheads indicate anticlinal cell walls. Scale bars represent 10µm. **(D)** Quantifications of the mean fluorescence intensity along anticlinal membranes of xylem-pole pericycle cells in the young maturation zone of 9-day old *PIN7:PIN7-GFP* seedlings, grown in the absence or presence of 1µM GLV10p. Fluorescence intensities are expressed in arbitrary units (a.u.). Individual datapoints (one per anticlinal membrane) and mean ± SD are shown. Statistical significance was determined via a Mann-Whitney test.

## Discussion

Previous research established the GLV6 and GLV10 signaling peptides as redundantly acting inhibitors of LR initiation (Fernandez *et al*., 2015, 2020). Knocking out both *GLV6* and *GLV10* leads to an increase in LR initiation events and an aberrant spacing of LR primordia, while an excess of GLV6 or GLV10 peptides results in a strong decrease in LR primordium density. The latter inhibitory effect was found to be associated with the occurrence of ectopic, and relatively symmetric, anticlinal cell divisions throughout the pericycle (Meng *et al*., 2012; Fernandez *et al*., 2015; Jourquin *et al*., 2022). This indicates that these peptides do not merely inhibit LRFC divisions, but somehow stimulate non-formative pericycle cell divisions. In agreement with this, we found that GLV6/10 peptides do not directly inhibit the expression of key regulators of LR initiation, but actually trigger an auxin-like response in their target cells (i.e. pericycle cells, including LRFCs). Our data indicate that this partial auxin response and the ectopic pericycle cell divisions do not rely on auxin perception or signaling. As the transcription of the *GLV6* and *GLV10* genes themselves is auxin inducible (Fernandez *et al*., 2020; Jourquin *et al*., 2022), this suggests that these peptides aid in the regulation of a subset of auxin response genes, downstream of the canonical auxin signaling pathway, which includes several cell cycle genes. This GLV-induced partial auxin response is thus likely the cause of the ectopic anticlinal pericycle cell divisions observed when GLV6/10 peptides are in excess, but does not readily explain how these peptides inhibit LR initiation. Interestingly, despite this auxin-like transcriptional response, we found that auxin itself fails to accumulate in LRFCs upon *GLV6*^*OE*^ or GLV10p treatment. Hence, we propose that the inhibitory effect of GLV6/10 signaling on LR initiation is caused by the absence of sufficiently high auxin levels and the associated lack of a full transcriptional auxin response in LRFCs.

Initially, we investigated whether the GLV6/10-induced inhibition of pre-branch site establishment might result from an effect on the conjugation of auxin to amino acids, since several group II *GH3* genes were differentially regulated upon *GLV6*^OE^. We hypothesized that GLV6/10 peptide signaling might stimulate IAA-AA conjugation via these GH3 enzymes, thereby reducing the levels of free auxin in LRFCs. In agreement with this, overexpression and dominant gain-of-function mutants of several *GH3* genes were reported to show strong reductions in emerged LR densities, while *gh3*.*123456* sextuple mutants were found to have an increased LR density, phenotypes that correspond to those of *GLV6*^*OE*^ lines and *glv6glv10* double mutants, respectively (Nakazawa *et al*., 2001; Takase *et al*., 2004; Xuan *et al*., 2015; Porco *et al*., 2016). Furthermore, *gh3*.*123456* mutants were found to be somewhat resistant to the effects of GLV10p treatments on auxin distribution and LR initiation. However, despite the clear disappearance of DR5:LUC-marked pre-branch sites, the levels of free IAA and its detectable conjugates and degradation products were found to be unaffected in root bends of *GLV6* overexpressing seedlings. These data thus indicate that *GLV6*^*OE*^ does not affect overall IAA levels in the root, but rather prevents its accumulation in LRFCs. This is likely the result of an effect on auxin transport, rather than a GH3-mediated inactivation of IAA. The GLV6-induced downregulation of *GH3*.*1, GH3*.*2, GH3*.*3* and *GH3*.*5* expression, which drew our attention to these genes in the first place, might actually be a result of the reduced auxin levels in pre-branch sites, rather than the cause. Transcription of these *GH3* genes is known to respond extremely quickly to even relatively small changes in auxin concentrations (Paponov *et al*., 2008). This could explain why these *GH3s* are among the very limited number of genes that show opposing responses to *GLV6*^*OE*^ and auxin treatment. The reason for the hyposensitivity of *gh3*.*123456* mutants to GLV10p treatment therefore remains speculative, but might result from the generally higher IAA concentrations in these mutants (Porco *et al*., 2016).

Since free auxin levels seemed to be unaffected by GLV6/10 peptide signaling, we investigated whether these peptides might influence LR initiation via an effect on auxin transport. Indeed, mutants in members of several classes of auxin transport proteins showed aberrant responses to GLV10p treatment, pointing to a connection between GLV6/10 signaling and auxin transport. The *ABCB1, ABCB4, ABCB19* and *ABCB21* auxin exporters are all known to be implicated in LR development (Noh *et al*., 2001; Lin and Wang, 2005; Santelia *et al*., 2005; Wu *et al*., 2007; Jenness *et al*., 2019), but due to the extremely stunted primary root growth of GLV10p-treated *abcb1abcb19* mutants and the lack of GLV10p-induced changes in ABCB1-GFP and ABCB19-GFP expression patterns, it remains unclear whether and how these ABCBs would be involved GLV6/10-RGI pathway. It is possible that GLV6/10 peptides affect auxin transport independently of ABCBs, but that the loss of ABCB1 and ABCB19, combined with the effect of superfluous GLV6/10 signaling, disorganizes the distribution of auxin to such an extent that root development is severely disrupted. It is in any case clear that *abcb1abcb19* mutants are not hyposensitive to GLV10p treatment, suggesting that these ABCBs are not strictly required for the response to GLV6/10 peptides. Similarly, the increased responsivity of *aux1* mutants to GLV6/10-induced inhibition of LR initiation suggests that AUX1 activity is not needed for GLV6/10-RGI signaling, but rather counters its effect. This is indeed plausible, as AUX1-mediated auxin import into LRFCs is known to enhance local auxin accumulation, which likely opposes the effect of GLV6/10 peptides (Marchant *et al*., 2002; Laskowski *et al*., 2008). The misexpression of AUX1 throughout large stretches of the pericycle is therefore unlikely to be the cause of the GLV10p-induced LR initiation defects. The PIN3 and PIN7 auxin export proteins on the other hand, seem to be imperative for the effect of GLV6/10 signaling on LR initiation, since *pin3pin7* double mutants displayed on almost complete resistance to GLV10p treatment. This behavior was not observed in *pin3* and *pin7* single mutants, in agreement with previous reports showing that both of these auxin exporters can act redundantly in several processes (Kleine-Vehn *et al*., 2010; Lewis *et al*., 2011; Marhavý *et al*., 2013; Wang *et al*., 2015). The inhibitory effect of GLV6/10 signaling on the establishment of auxin maxima in LRFCs is thus likely achieved via the PIN3 and PIN7 auxin exporters. GLV10p treatment did not trigger any apparent changes in *PIN3:PIN3-GFP* expression, but resulted in stronger expression of *PIN7:PIN7-GFP* throughout the pericycle, with significantly increased PIN7-GFP levels in anticlinal membranes of xylem-pole pericycle cells. GLV6/10 signaling might thus stimulate auxin export from its target cells by increasing PIN7 abundance. In agreement with this, ethylene-induced increases in *PIN3* and *PIN7* expression were previously shown to result in an inhibition of LR initiation (Lewis *et al*., 2011), and *pin3pin7* double mutants are known to show defects in LR spacing (Benková *et al*., 2003; Laskowski *et al*., 2008). Additionally, previous research has indicated that GLV peptide signaling can enhance PIN2 levels in the plasma membrane of epidermal and cortical cells in the root apical meristem, which in turn affects gravitropic root growth (Whitford *et al*., 2012). Although our data do not point to the involvement of PIN2 in the GLV6/10-RGI pathway during LR initiation, the modulation PIN abundance in the plasma membrane via GLV signaling might be a general mechanism. The effect of GLV6/10 signaling on PIN7 abundance might be at least partially transcriptional, since *PIN7* expression was found to be significantly upregulated upon *GLV6*^OE^, albeit with a very low fold change (FC=1.2) (Supplementary table 1). However, the possibility of post-translational effects of GLV6/10 signaling on PIN3 and PIN7 abundance and activity cannot be excluded.

Although further research is required to elucidate the exact molecular mechanism via which GLV6/10 peptides affect PIN plasma membrane abundance and auxin transport, the results presented here allow us to formulate a model of the way in which the interplay between auxin and GLV6/10-RGI signaling might regulate the spacing of LRs and their development (Fig. 7). LRFCs are believed to be primed for LR formation by recurrent peaks of auxin signaling activity in the elongation zone, the frequency of which largely determines the distance between consecutive LRs (De Smet et al., 2007; Moreno-Risueno et al., 2010; Xuan et al., 2015). However, it has been shown that, even in wild-type seedlings, multiple pairs of LRFCs are often specified in close proximity to one another, but that typically only one pair will give rise to a primordium, indicating that the others are somehow silenced (Toyokura *et al*., 2019). The TARGET OF LBD SIXTEEN 2 (TOLS2) peptide and its receptor RLK7 were found to be involved in this process, and we believe that GLV6/10-RGI signaling plays a similar role. Auxin is well known for its ability to regulate its own distribution via feedback loops on its biosynthesis and degradation as well as its intercellular transport, typically generating well defined and regularly spaced auxin maxima in both the shoot and the root (Mironova *et al*., 2010; del Bianco and Kepinski, 2011; Sassi and Vernoux, 2013). We propose that the GLV6/10 signaling pathway is an important component of this self-organizing capacity during the establishment of pre-branch sites along the primary root. When GLV6/10 peptides are in overabundance, PIN7 levels are increased throughout the xylem-pole pericycle. The increased abundance of this auxin exporter prohibits the accumulation of auxin in xylem-pole pericycle cells, including LRFCs, thereby preventing the establishment of the auxin response maxima required for LR initiation. Additionally, an excess of GLV6/10 peptides can trigger ectopic anticlinal pericycle cell divisions in an auxin independent manner. In natural conditions, however, the accumulation of auxin in LRFCs precedes and indeed triggers *GLV6/10* expression (Fernandez *et al*., 2020; Jourquin *et al*., 2022). In these circumstances, local auxin levels are likely sufficiently high to maintain further auxin accumulation, despite the GLV-induced stimulation of auxin export. Endogenous GLV6/10 peptides are therefore unable to inhibit LR initiation in the founder cells from which they are secreted (Fig. 7). Diffusion of GLV6/10 peptides through the apoplast might create a zone around these LRFCs (or a developing primordium) where peptide levels are sufficiently high to enhance PIN7 abundance and inhibit the accumulation of auxin, thereby effectively preventing LR initiation in neighboring pericycle cells (Fig. 7). The balance between auxin and GLV6/10 peptide levels and their spatiotemporal distributions along the pericycle might thus create inhibitory zones around primordia in which the peptides prevent the initiation of LR primordia in close proximity to pre-existing ones. Additionally, at the level of the primordium itself, the stimulatory effect of GLV6/10 peptides on pericycle cell divisions could potentially contribute to primordium development in concert with auxin. In this model, GLV6/10 peptide signaling thus serves as an auxin inducible system that contributes to the robust patterning of auxin response maxima along the primary root by stimulating auxin export from XPP cells via PIN3 and PIN7.

**Fig. 7.**
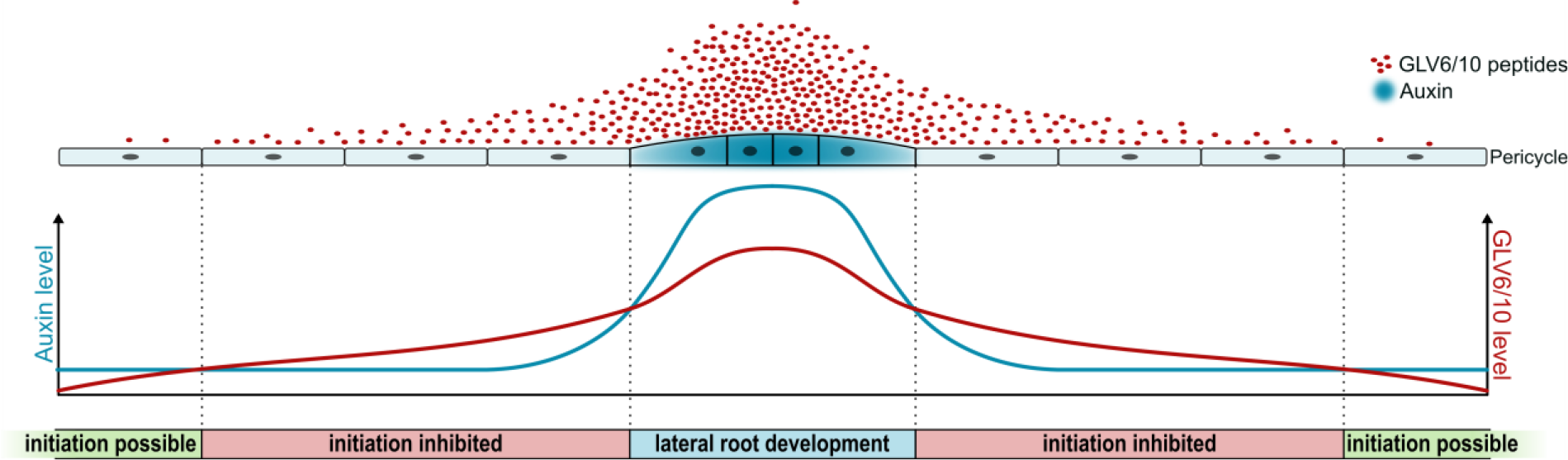
Proposed model of the role of GLV6/10 signaling during LR initiation in relation to auxin. Model for the regulation of LR initiation and spacing via the balance between auxin and GLV6/10 peptide levels. The accumulation of auxin in LRFCs induces the expression of *GLV6* and *GLV10*, resulting in the secretion of GLV6/10 peptides into the apoplast. Here they diffuse outward, creating a concentration gradient around the initial pair of LRFCs. Whitin these LRFCs, the high auxin/GLV ratio permits further accumulation of auxin, and the effect of GLV6/10 peptides on pericycle cell divisions might even aid in the correct patterning and/or the progress of the initial LRFC divisions. Around the initiation site, a zone is created in which the low auxin/GLV ratio inhibits the accumulation of auxin in pericycle cells, most probably via the GLV-induced increase in PIN7 abundance. As a result, the initiation of new LR primordia within this zone is inhibited. Further outward, this inhibitory zone is flanked by zones in which GLV6/10p levels are sufficiently low to allow for LR initiation.

## Acknowledgments

We thank Dr. Angus Murphy for providing us with the *abcb* mutant seeds used in this study, and Dr. Veronique Storme for her advice on statistical analyses.

## Author Contributions

Conceptualization, J.J., A.I.F., and T.B.; Methodology, J.J., J.S., K.L., A.I.F., and T.B.; Formal Analysis, J.J.; Investigation, J.J., K.X., and J.S.; Resources, K.L., and T.B.; Writing-Original Draft, J.J.; Writing – Review & Editing, J.J., A.I.F., and T.B.; Visualization, J.J.; Supervision, J.J., A.I.F., and T.B.; Funding Acquisition, J.J., A.I.F., and T.B.

## Data Availability

All data supporting the findings of this study are available within the paper and within its supplementary materials published online.

## Supplementary Figures

**(S1).**
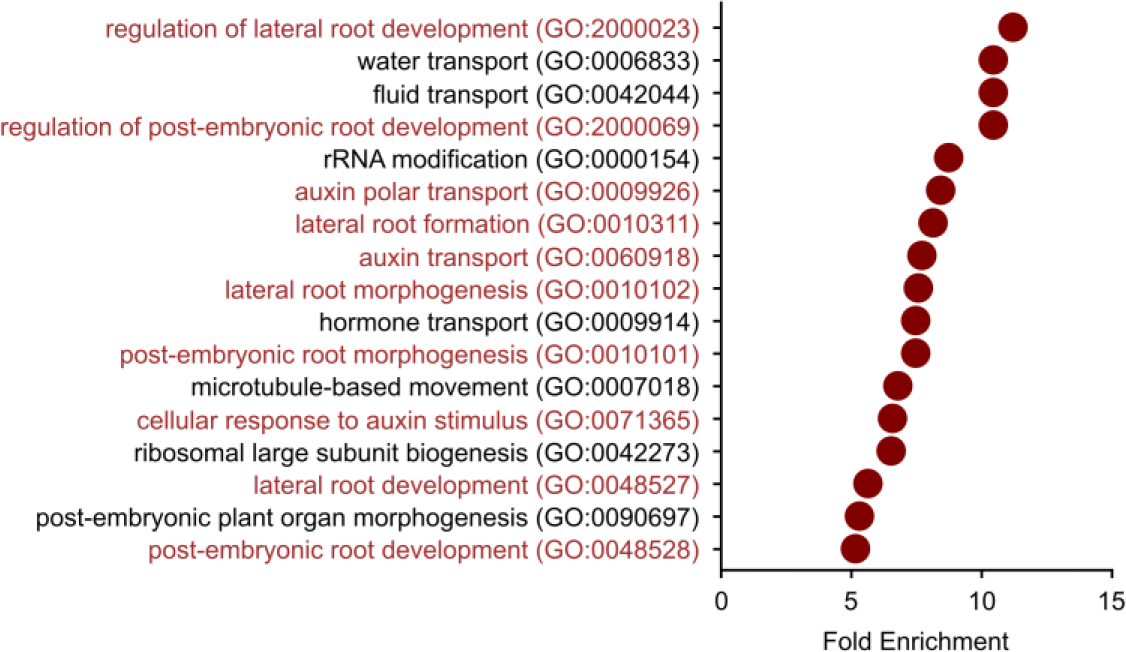
Gene Ontology enrichment analysis on all GLV6 response genes identified in the *GLV6*^OE^ RNA-sequencing experiment (Jourquin *et al*., 2022). All significantly enriched GO terms (adj. p < 0.05) with a fold enrichment higher than 5 are shown. Statistical significance was determined via a Fisher’s Exact test with Bonferroni correction. GO terms associated with LR development and auxin signaling or transport are highlighted in red.

**(S2).**
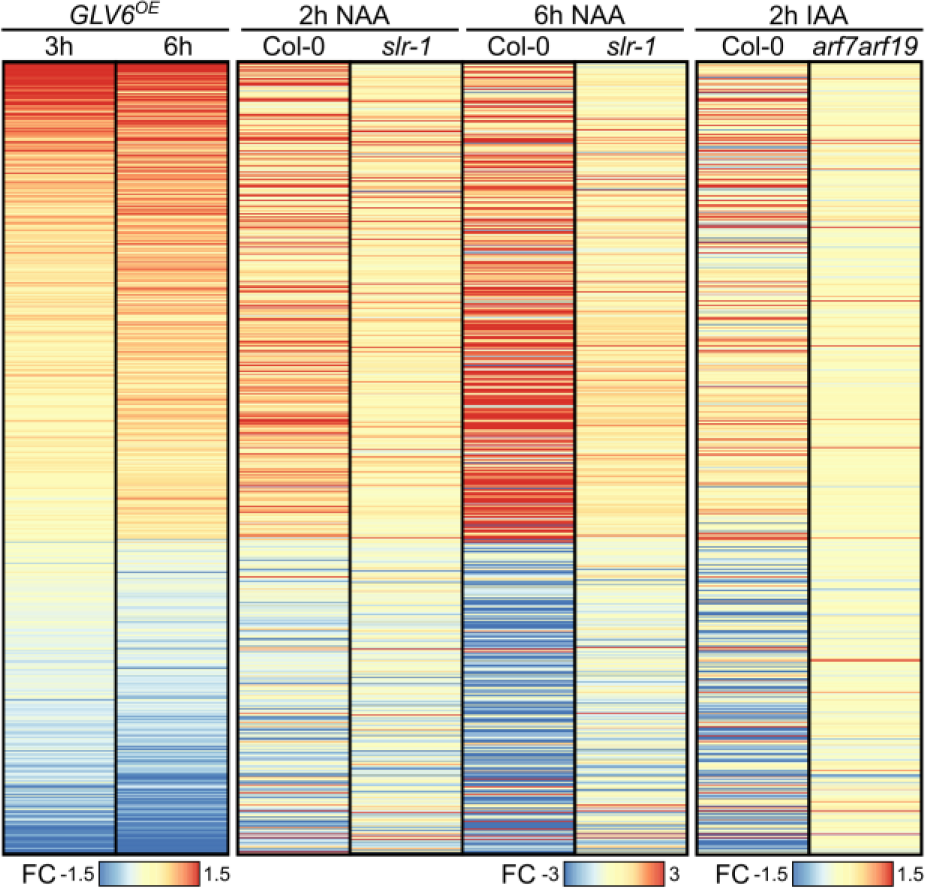
Heatmap depicting the fold changes (FC) of the 685 genes that show a significant response to GLV6 overexpression as well as NAA or IAA treatment as shown by *jourquin et al*., *2022; Vanneste et al*., *2005 and Okushima et al*., *2005* respectively. A list of these genes and their exact transcript fold changes can be found in *supplementary table 1*.

**(S3).**
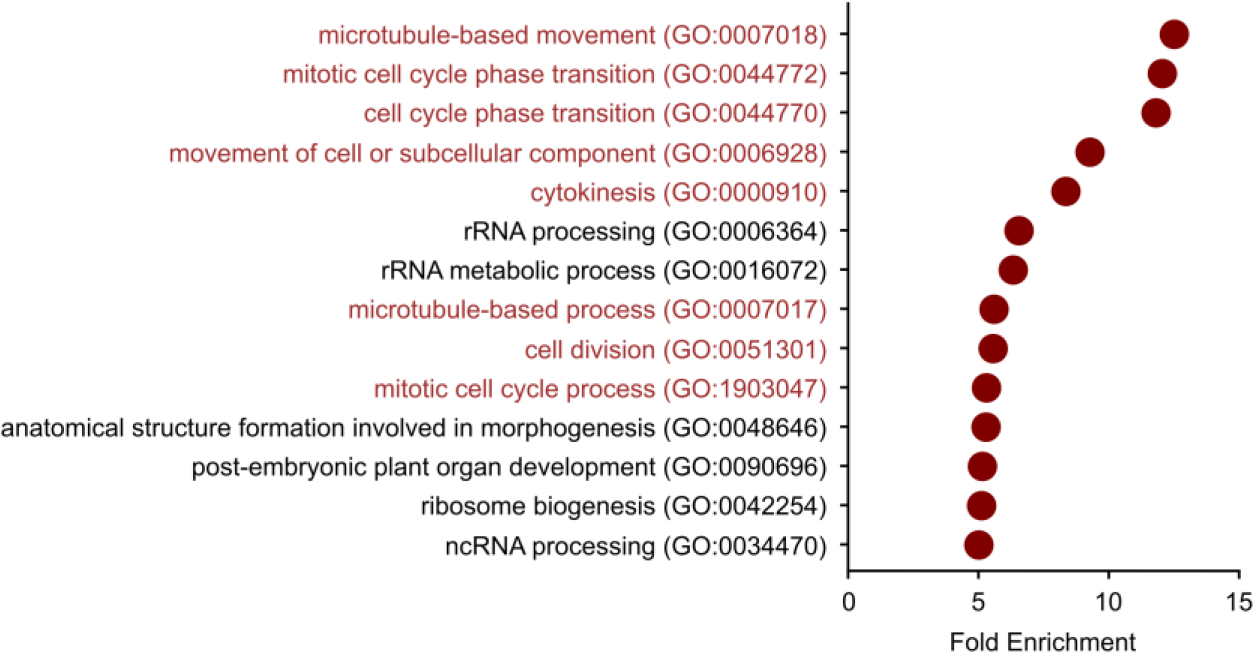
Gene Ontology enrichment analysis on “late” GLV6 response genes identified in the *GLV6*^OE^ RNA-sequencing experiment as genes that are differentially regulated 6 hours after induction of *GLV6* overexpression (Jourquin *et al*., 2022). All significantly enriched GO terms (adj. p < 0.05) with a fold enrichment higher than 5 are shown. Statistical significance was determined via a Fisher’s Exact test with Bonferroni correction. GO terms associated with cell division are highlighted in red.

**(S4).**
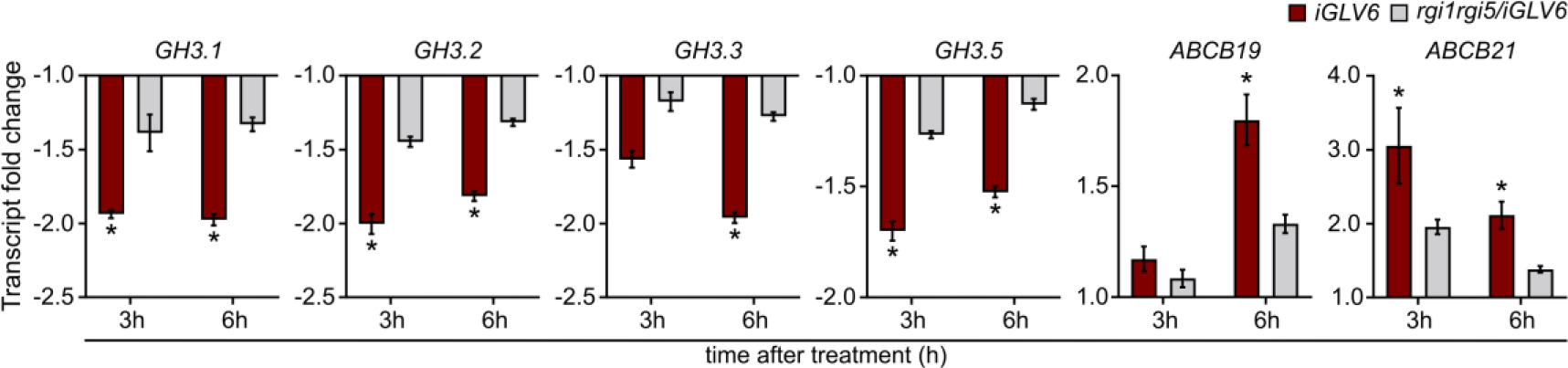
Fold changes in the expression levels of *GH3*.*1, GH3*.*2, GH3*.*3, GH3*.*5, ABCB19* and *ABCB21*, 3h and 6h after estradiol-induced *GLV6* overexpression in root bends of *iGLV6* and *rgi1rgi5/iGLV6* seedlings, as determined via the *GLV6* overexpression RNA-sequencing experiment (Jourquin *et al*., 2022). Significant fold changes are indicated with an asterisk (adj. p-value < 0.05), n=3. Statistical significance was determined via a Wald-test with Benjamini-Hochberg correction.

**(S5).**
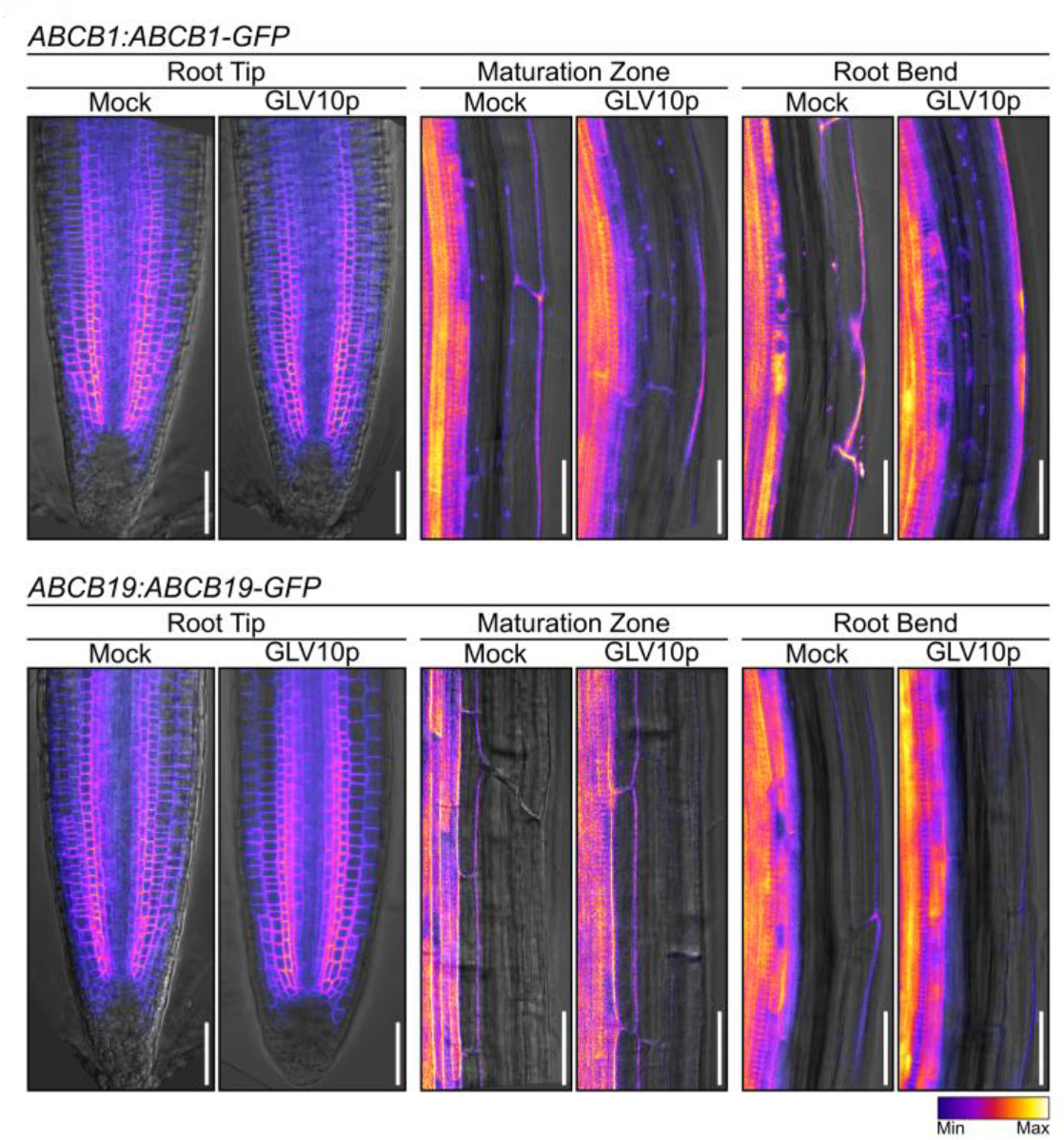
Primary root lengths of 12-day old wild-type seedlings and several auxin transport mutants, treated or not with 100nM GLV10p. Mean root length ± SE are shown. For each genotype, the percentual in-or decrease in primary root length as a result of GLV10p treatment is indicated. Asterisks indicate an in- or decrease that is significantly different (p-value < 0.05) from that observed in the wild-type control (none in this case). Statistical significance was determined via a two-way ANOVA and a post-hoc Dunnett’s test, n=15.

**(S6).**
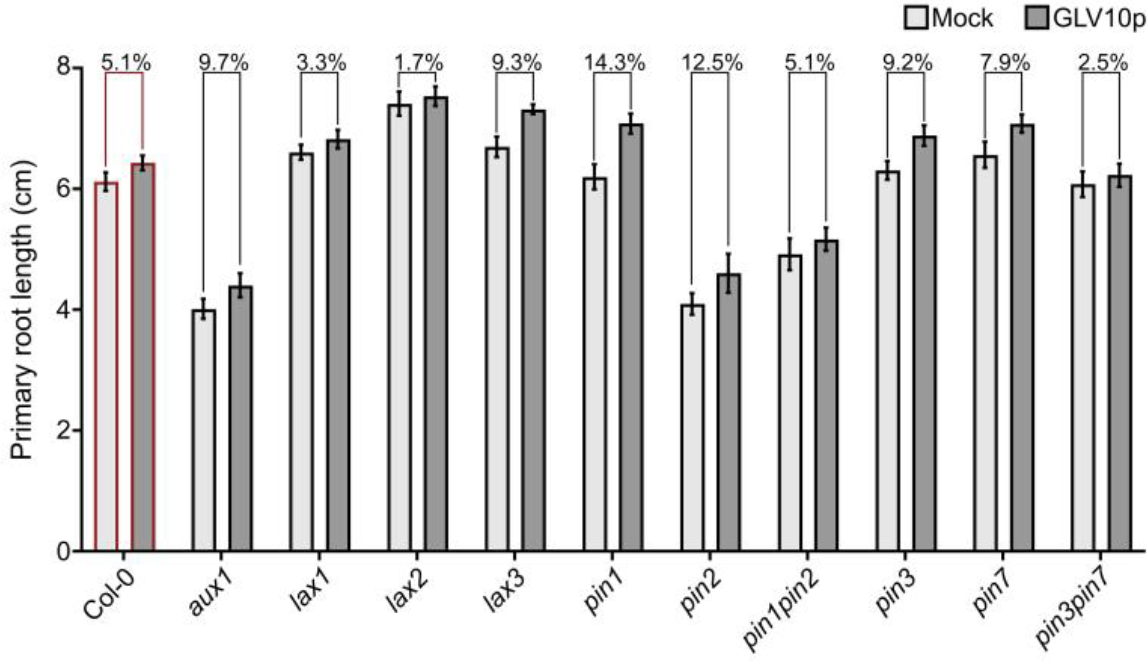
Confocal images of *ABCB1:ABCB1-GFP* and *ABCB19:ABCB19-GFP* expression in gravistimulated roots that were treated or not with 1µM GLV10p at 8h after gravistimulation and imaged 18h after gravistimulation. Images were taken from the root tips (left panels), sections of the maturation zone above the root bends (middle panels), and sections within the root bends (right panels). Scale bars represent 30µm.

